# *Boechera* or not? Genomic insights and taxonomic reassessment of the misclassified Asian species *B. calcarea* (Brassicaceae)

**DOI:** 10.1101/2025.01.28.635196

**Authors:** Terezie Mandáková, Milan Pouch, Petra Hloušková, Dmitry A. German, Michael D. Windham, Pavel Trávníček, Martin A. Lysak

**Author notes:** For correspondence: TM and MAL.

## Abstract

The genus *Boechera* (Brassicaceae) serves as a model system for studying apomictic reproduction and ecological adaptations, with most species concentrated in North America. Rare occurrences of *Boechera* species beyond their typical range offers unique opportunities to investigate genome evolution in extralimital environments. One such species, *B. calcarea*, was described from the Chandalaz Range in northeastern Asia (Russia). This study aimed to investigate the genome structure and evolutionary history of *B. calcarea*. However, our analyses reveal that the species does not belong to *Boechera*. Instead, an integrative approach combining cytogenetic, phylogenetic and repeatome analyses identified the species as a member of an ancestral clades within the tribe Arabideae. The diploid *Parryodes calcarea* (Dudkin) D.A.German & Lysak (2*n* = 16) exhibits Arabideae-specific chromosomal signatures, including multiple centromere repositionings. These findings clarify the misattribution of *P. calcarea* to *Boechera*, leaving *Boechera falcata* and *Borodinia macrophylla* as the only Old World representatives of the Boechereae. This study highlights the importance of integrative methodologies in resolving taxonomic ambiguities and provides new insights into the diversification of the largest cruciferous tribe, Arabideae.

## INTRODUCTION

The classification of plants into genera and species is fundamental to botany, offering a framework for understanding biodiversity, ecological adaptations, and the evolutionary history of life on Earth. Traditionally, plant taxonomy relied heavily on morphological characteristics, including fruit shape, trichome morphology, and reproductive traits, to group species into genera. However, these phenotypic markers, while practical, often fail to reflect true evolutionary relationships. Convergent evolution can obscure phylogenetic signals, leading to erroneous classifications (Smith et al., 2020). Recent advancements in molecular phylogenetics and cytogenetics have revolutionized the field, providing botanists with powerful tools to unravel evolutionary histories and reassess taxonomic boundaries, especially in plant families with frequent morphological convergences (Mandáková et al., 2019; Guo et al., 2021; Mandáková and Lysak, 2022; Hendriks et al., 2023).

The family Brassicaceae, comprising approx. 4,060 species in 372 genera, ranks among the largest and most ecologically diverse plant families (Hendriks et al., 2023). Its species exhibit remarkable morphological, physiological, and ecological variability, which contributes to the taxonomic complexity of the family. Within Brassicaceae, the tribe Arabideae has presented persistent challenges due to its morphological overlap and polyphyly in earlier classifications. Historically, classifications relied on traits such as latiseptate siliques, branched trichomes and accumbent cotyledons to group morphologically similar species under a single taxonomic framework (Hopkins, 1937; Rollins, 1941). However, molecular and cytogenetic studies revealed that the traditional circumscription of *Arabis* was polyphyletic, encompassing multiple distinct evolutionary lineages. These findings necessitated numerous transfers of former Arabideae species to genera outside the tribe, such as *Arabidopsis, Boechera, Catolobus, Pennellia*, and *Turritis* (Koch et al., 1999, 2001, 2003, 2012; Al-Shehbaz, 2003, 2005; Warwick and Souder, 2005; Al-Shehbaz et al., 2006; Kiefer et al., 2009; Hendriks et al., 2023).

The genus *Boechera*, segregated from *Arabis* by Löve and Löve (1976), has since been confirmed as a monophyletic group based on molecular and cytogenetic evidence (Al-Shehbaz, 2003). *Boechera* is distinguished by its base chromosome number of *x* = 7, in contrast to *x* = 8 in *Arabis s. str*., as well as by its unique reproductive biology, including the production of asexual seeds via diverse developmental pathways (Carman et al., 2019) and the widespread occurrence of diploid apomixis (Al-Shehbaz, 2003; Koch et al., 2003; Beck et al., 2012; Mandáková et al., 2020a). Morphologically, *Boechera* is characterized by dendritic, irregularly bifurcate or sessile leaf trichomes, often bent and/or pedent fruits and some other features not or rarely found in *Arabis* (Al-Shehbaz 2003). These traits, along with molecular and cytological evidence, clearly distinguish the two genera.

While *Boechera* primarily occurs in North America, particularly in the western United States, the discovery of species outside the continent provides valuable insights into the evolutionary history and dispersal mechanisms of the genus. These extralimital taxa challenge the notion of *Boechera* as a strictly North American lineage and suggest historical biogeographic connections between the continents (Al-Shehbaz, 2003; Windham and Al-Shehbaz, 2006, 2007). Such species may have arisen through long-distance dispersal events or as remnants of a broader ancestral distribution facilitated by climatic and geological phenomena, such as the Bering Land Bridge during glacial periods (Abbott et al., 2000; Brigham-Grette et al., 2013). These events likely allowed cold-adapted Brassicaceae, including ancestral *Boechera*, to migrate between North America and Asia, leading to isolated populations that evolved into distinct species (Warwick and Sauder, 2005; Koch et al., 2012).

The most extensively studied extralimital Boechereae species is *B. falcata*, found in northeastern Asia, including Siberia and the Russian Far East. Morphological similarity of *B. falcata* with its North American relatives (e. g., with *B. holboellii*, the type of *Boechera*: Busch, 1926; Schulz, 1936; Berkutenko, 1983, in all cases under *Arabis*) was noticed long ago, and its inclusion in the genus was subsequently supported by phylogenetic studies (Kiefer et al., 2009, 2014). Cytogenetical analysis revealed a diploid chromosome number of 2*n* = 14, consistent with the genus *Boechera* and further corroborating its phylogenetic placement (Yurtsev and Zhukova, 1972; Berkutenko and Gurzenkov, 1976). Another species of particular interest is *B. calcarea*, described by Doudkin and Volkova (2013) from the Chandalaz Range in Primorsky Territory, Russia. The Chandalaz Range, described as the largest calcareous massif in the Russian Far East, was suggested to serve as a refugium for numerous rare plant species (Doudkin, 1998). According to Doudkin and Volkova (2013), *B. calcarea* was characterized as an obligate calciphile thriving in nutrient-poor, high-calcium soils and morphologically was distinguished from *B. falcata* by larger fruits, forked (vs. substellate) trichomes, and pale lavender (vs. pink) petals (**Suppl. Fig. 1A**). *Boechera calcarea* was reported to exhibit a diploid chromosome number of 2*n* = 14, consistent with the dominant chromosome number in *Boechera* (Doudkin and Volkova, 2013; **Suppl. Fig. 1B**). Unlike *B. falcata, B. calcarea* has not been extensively studied in terms of its genetic structure or reproductive biology, leaving significant gaps in understanding its evolutionary history.

**Fig. 1.**
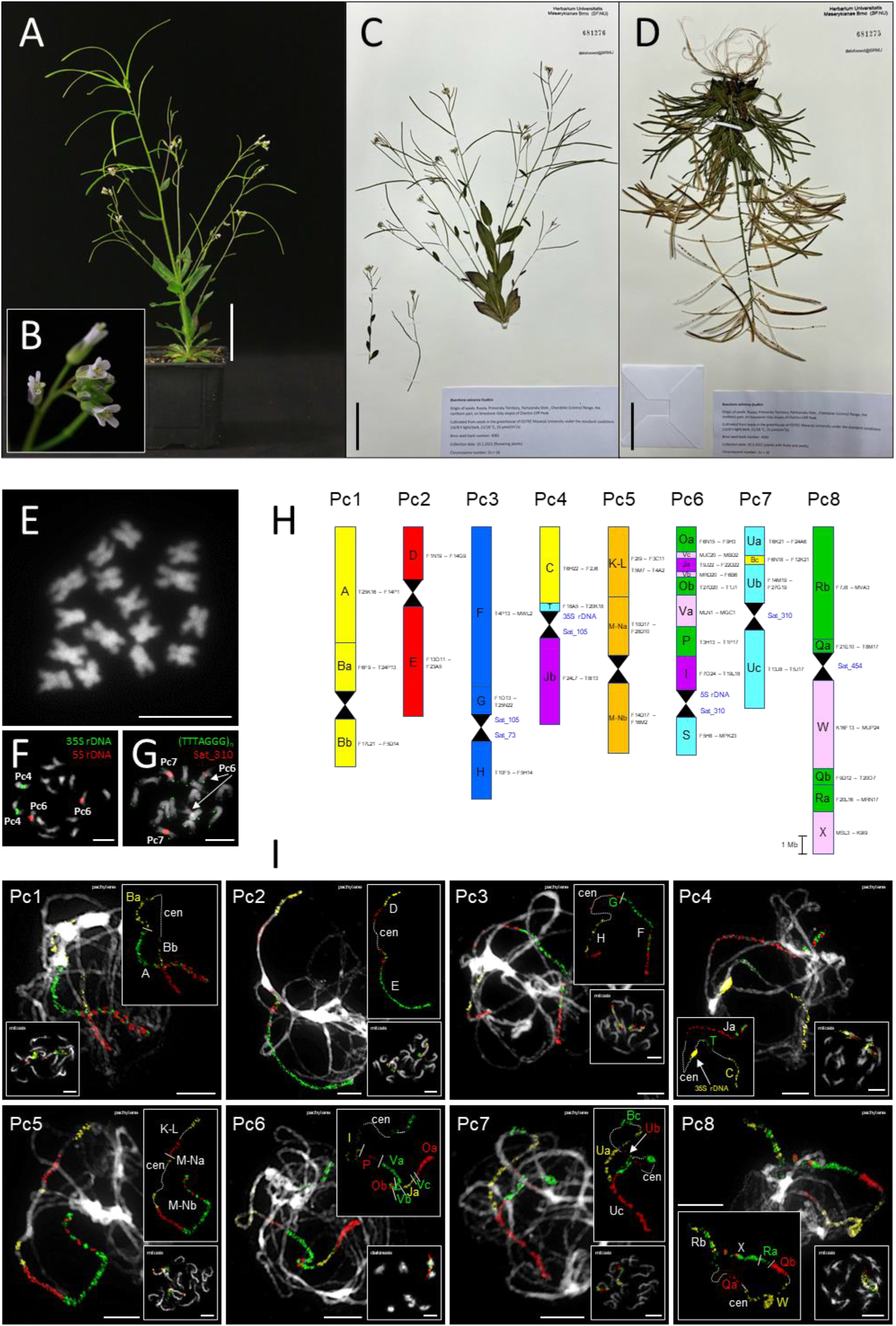
Genome structure of *Parryodes calcarea*. **(A)** Photograph of cultivated plant. **(B)** Close-up of inflorescence. **(C)** Herbarium specimen of flowering plant accession investigated, deposited in the herbarium of Masaryk University (BRNU). **(D)** Herbarium specimen of plant bearing fruits. **(E)** Mitotic chromosome spread prepared from a root tip, showing 2*n* = 16 chromosomes. (F) Mitotic chromosomes hybridized with 35S and 5S rDNA probes. **(G)** Mitotic chromosomes hybridized with telomeric and centromeric satellite Sat_310 probes. **(H)** Karyotype structure inferred from comparative chromosome painting (CCP), with the eight individual chromosomes labeled as Pc1–Pc8. Capital letters (A to X) represent the 22 genomic blocks (GBs) of the Ancestral Crucifer Karyotype (ACK; Mandáková et al., 2019), some of which are reshuffled into sub-blocks (e.g., Ba, Bb, Bc). Each GB is color-coded to correspond to its original position on the eight chromosomes of the ACK. Hourglass symbols indicate centromeres, and Arabidopsis BAC clones are provided as markers for each (sub-)block. Chromosomes are drawn to scale (scale bar = 1 Mb). **(I)** The eight chromosomes (Pc1–Pc8) were visualized through CCP using Arabidopsis BAC contigs as painting probes applied to pachytene, mitotic, and diakinesis chromosome spreads. Painting signals are shown in experimental colors, reflecting the fluorochromes used to label specific GBs. When a GB is labeled with a single fluorochrome, its letter is displayed in the corresponding color. For subdivided GBs labeled with multiple fluorescent dyes to indicate their orientation, the letters are shown in white. Centromeres are marked as “cen”. All chromosomes were counterstained with DAPI and presented as grayscale images. Scale bars are 5 cm for plant photographs (A, C, D) and 10 µm for chromosome images (E, F, G, I).

Here, we present a detailed genomic characterization of *B. calcarea*. Surprisingly, our analyses revealed that the species does not belong to *Boechera* or any other genus of the tribe Boechereae. Instead, it represents a distinct species in the tribe Arabideae, recently reclassified, based on the results of the present study, as *Parryodes calcarea* (Dudkin) D.A.German & Lysak (Chepinoga et al., 2024). This leaves *B. falcat*a and *Borodinia macrophylla* as the only Boechereae species of the Old World.

## MATERIALS AND METHODS

### Plant material

Seeds were collected by Roman V. Doudkin on exactly the same locality from which *B. calcarea* was described (Doudkin and Volkova, 2013): Russia, Primorsky Territory, Partizansky Distr., Chandalaz (Lozovy) Range, the northern part, on limestone rocky slopes of Chertov Cliff Peak. Plants were cultivated from seeds in the greenhouse of CEITEC Masaryk University under the standard conditions (16/8 h light/dark, 21/18 °C, 15 µmol/m2/s) (**Fig. 1A, B**). Specimens are deposited at the Herbarium of Masaryk University (BRNU): BRNU sheet numbers: 681276 (flowering plant; **Fig. 1C**; https://brnu.jacq.org/BRNU681276) and 681275 (plant with fruits and seeds; **Fig. 1D**; https://brnu.jacq.org/BRNU681275).

### Chromosome preparation

Actively growing root tips were harvested from cultivated plants, pre-treated with ice-cold water for 12 hours, and subsequently fixed in a freshly prepared ethanol:acetic acid solution (3:1) for 24 hours at 4°C. The fixed samples were then stored at −20 °C for future use. Entire inflorescences were similarly fixed overnight in freshly prepared ethanol:acetic acid solution (3:1), then transferred and stored at-20 °C until processing. Mitotic and meiotic (pachytene) chromosome preparations were conducted following the protocol described by Mandáková and Lysak (2016a). Prior to analysis, suitable slides were pre-treated with RNase (100 mg/ml) and pepsin (0.1 mg/ml).

### Probe preparation

A total of 674 chromosome-specific BAC clones from *Arabidopsis thaliana* (Arabidopsis) were employed, grouped into contigs based on 22 genomic blocks as described by Mandáková et al. (2019). Probes were designed according to the known genomic structure of *Pseudoturritis turrita* and other Arabideae crown group species (Mandáková et al., 2020b; Nowak et al., 2020), reflecting the configuration of the eight chromosomes. These BAC contigs were used for chromosome painting. To achieve detailed chromosome structure characterization and precise localization of centromeres, initial experiments were followed by fine-scale painting using shorter BAC (sub)contigs. Individual, differentially labeled BAC clones were applied for BAC-by-BAC centromere localization. The localization of 5S and 35S rDNA loci was achieved using clone pCT 4.2 (corresponding to the 500-bp 5S rRNA repeat, GenBank accession M65137) and Arabidopsis BAC clone T15P10 (GenBank accession AF167571), respectively. An Arabidopsis-like telomeric probe was prepared using the protocol outlined by Ijdo et al. (1991). Additionally, four species-specific tandem repeats (Sat_73, Sat_105, Sat_177, Sat_310, and Sat_454) were identified and prepared as described below. All probes were labeled with biotin-dUTP, digoxigenin-dUTP, and Cy3-dUTP via nick translation, then pooled, precipitated, and resuspended in 20 μl of hybridization mixture (50% formamide and 10% dextran sulfate in 2× SSC) per slide, following the methodology of Mandáková and Lysak (2016b).

### Comparative chromosome painting

Chromosome and probe denaturation were performed simultaneously on a hot plate at 80 °C for 2 minutes, followed by overnight incubation in a humidified chamber at 37 °C. Post-hybridization washes were carried out in 20% formamide in 2× SSC at 42 °C. Fluorescent detection was conducted according to Mandáková and Lysak (2016b) as follows: biotin-dUTP was detected using avidin–Texas red, with signal amplification by goat anti-avidin–biotin and avidin–Texas red; digoxigenin-dUTP was detected using mouse anti-digoxigenin followed by goat anti-mouse Alexa Fluor 488. Chromosomes were counterstained with 4′,6-diamidino-2-phenylindole (DAPI; 2 μg/ml) in Vectashield.

### Image processing

Fluorescent signals were analyzed and photographed using a Axioimager epifluorescence microscope (Zeiss) and a CoolCube camera (MetaSystems). Images were acquired individually for the four fluorochromes using corresponding excitation and emission filters (AHF Analysentechnik). The captured images were pseudocolored and merged using Adobe Photoshop CS6 software (Adobe Systems). Circular visualizations of chromosome-scale pseudomolecules were prepared using Circos (Krzywinski et al., 2009).

### Phylogenetic analysis

Genomic DNA was extracted from young leaves using the NucleoSpin Plant II kit (Macherey-Nagel). Internal transcribed spacers (ITS1 and ITS2) were amplified using the primer pair ITS1/ITS4 (White et al., 1990), and the purified amplicons were Sanger sequenced at Macrogen. The resulting ITS sequences were deposited in GenBank (**Suppl. Tab. 1**). These sequences were then analyzed using BrassiBase (Kiefer et al., 2014), where a Maximum Likelihood (ML) tree was inferred using the “Phylogenetics Tool”. The resulting tree indicated that investigated accession clusters within the clade of the Arabideae tribe. To enhance the analysis, ITS sequences of all available representative of the Arabideae were retrieved from BrassiBase, aligned with MAFFT v7.490 using default parameters (Katoh and Standley, 2013), and manually trimmed. The final ML tree was generated using IQ-TREE v2.2.0 with 10,000 ultrafast bootstrap (UFbootstrap) replicates (Minh et al., 2020). Nodes with UFbootstrap support below 50 were collapsed using Newick Utilities v1.6.0 (Junier and Zdobnov, 2010). As IQ-TREE produces unrooted trees, *Pseudoturritis turrita*, a basal taxon of the Arabideae tribe, was designated as the outgroup, and the tree was subsequently rerooted.

Low-coverage genome sequencing of the studied species was performed using the Illumina NovaSeq 6000 platform with 250 bp paired-end reads. Adapter sequences and low-quality reads were filtered out using Trimmomatic v0.39 (Bolger et al., 2014) with the following parameters: ILLUMINACLIP:TruSeq3-PE.fa:2:30:10:2:true, LEADING:10, TRAILING:10, SLIDINGWINDOW:4:20, and MINLEN:40. De novo assembly of the plastome was conducted with GetOrganelle v1.7.7.0 (Jin et al., 2020) using parameters -w 185,-R 15,-k 21,45,65,85,105,127, and the *Scapiarabis saxicola* plastome (MK637807) as the seed. Plastome annotation was performed using the GeSeq v2.03 pipeline (Tillich et al., 2017), and a graphical visualization was generated with OGDRAW v1.3.1 (Greiner et al., 2019). The annotated plastome was submitted to GenBank (**Suppl. Tab. 1**).

To confirm the phylogenetic placement of the studied species within the Arabideae tribe, the plastid trnL-trnF region was analyzed. Sequences for this region were sourced from the study by Karl and Koch (2013), and multiple sequence alignments along with ML tree inference were performed using the same methods as for the ITS region. Furthermore, the same phylogenetic analysis pipeline was applied to the available complete plastomes of Arabideae species (Suppl. Tab. 1), which were either downloaded from GenBank or obtained from the study by Hendriks et al. (2023). One copy of the inverted repeats (IRs) was removed from the plastome alignment, which was generated using MAFFT, manually reviewed, and trimmed. The final ML tree was inferred using the previously described methods and further refined with phytools v2.3-0 (Revell, 2024).

### Genome size estimation

The genome size was estimated by flow cytometry. The suspension of isolated nuclei was prepared from fully developed intact leaves according to Doležel et al. (2007) and treated by propidium iodide and RNAase IIA (both 50 μg/ml) at room temperature for 5 min and analyzed using a Partec CyFlow cytometer. A fluorescence intensity of at least 5,000 particles was recorded. *Carex acutiformis* (1C = 432.8 Mb; Chumová et al., 2021) served as the primary reference (internal) standard.

### Repeat identification and annotation

Low-coverage genome sequencing of the studied species was performed using the Illumina NovaSeq 6000 platform, generating 250 bp paired-end reads. These data were utilized to assemble the plastid genome and characterize repetitive sequences. For clustering analysis, chloroplast reads were removed using Bowtie2 (v2.4.5), with the *Scapiarabis saxicola* chloroplast genome sequence (GenBank: MK637807.1) serving as the reference for read mapping. Non-chloroplast reads were filtered for quality, requiring a minimum quality score of 20% and at least 90% of nucleotides above this threshold. The reads were trimmed to a uniform length of 150 bp and interlaced. A subset of reads, representing 0.5× genome coverage (2,214,332 reads at 150 bp), was randomly sampled and used as input for the RepeatExplorer2 pipeline (Novák et al., 2020). Default clustering parameters were applied for species-specific analyses using 64 GB of memory. Superclusters were automatically annotated using the REXdb database (Neumann et al., 2019) and manually verified and corrected. Tandem repeats were identified using both RepeatExplorer2 and TAREAN (Novák et al., 2017). For *S. saxicola* (ERR4205135), individual clustering analysis was performed using reads pre-processed identically to those from the studied species. This resulted in 1,106,670 reads (100 bp each), which were input into the RepeatExplorer2 pipeline using default parameters.

To confirm the placement of the studied species within the Arabideae tribe, two comparative clustering analyses were conducted. The first utilized low-coverage Illumina sequencing data, while the second combined Illumina data with Hyb-Seq data. The dual approach accounted for the reduced representation of repetitive sequences in Hyb-Seq datasets.

Comparative analysis with Illumina data: This analysis included sequencing data from four Arabideae species (*Arabis auriculata, Ar. cypria, Aubrieta canescens*, and *Pseudoturritis turrita*; Mandáková et al., 2020b), along with publicly available Illumina reads for *S. saxicola* (ERR4205135). All reads were trimmed to 100 bp for consistency and sampled to approximately 0.1× genome coverage per species, resulting in 2,660,000 reads. For *S. saxicola*, genome size was unknown, so sampling followed the same parameters as for the studied species. Clustering was performed using RepeatExplorer2 with default settings.

#### Comparative analysis with Hyb-Seq data

The second analysis combined Illumina sequencing data for the studied species with Hyb-Seq data for *Parryodes axilliflora* (SRR22519503), *Scapiarabis saxicola* (SRR22519334) and *Acirostrum alaschanicum* (SRR22519361) (Hendriks et al., 2023). Reads corresponding to probe sets were filtered by mapping to the Brassicaceae bait sequences from Nikolov et al. (2019) and Angiosperms353 (Johnson et al., 2018) using Bowtie2 (v2.4.5). All reads were trimmed to 100 bp and randomly sampled to 500,000 reads per species (2,000,000 reads total). Clustering was performed in RepeatExplorer2 using default settings and the comparative clustering option.

Comparative clustering visualizations were generated within the RepeatExplorer2 Galaxy web interface (https://repeatexplorer-elixir.cerit-sc.cz/). Shared tandem repeats were aligned using MAFFT (v7.490, default settings) and manually adjusted in Geneious Prime 2023.0.1 (https://www.geneious.com).

### Oligoprobe design

In total, five tandem repeats with monomer lengths ranging from 73 to 454 bp were selected for further investigation (Sat_73, Sat_104, Sat_177, Sat_310, and Sat_454; **Suppl. Tab. 2**). Oligoprobes for these tandem repeats were designed based on reconstructed monomer consensus sequences generated using the TAREAN pipeline. The most conserved regions within the consensus sequences, featuring suitable GC content (30– 50%), were manually selected for 60-bp oligoprobes using the Geneious Prime 2023.0.1 software package (https://www.geneious.com). Regions with low potential for self-annealing and hairpin formation were prioritized during the design process. Unmodified synthetic single-stranded DNA (ssDNA) oligonucleotides were dissolved in ultrapure water to a final concentration of 100 μM. Equal molar amounts of complementary oligonucleotides were mixed in a 0.2-ml tube, heated to 95°C for 5 minutes in a PCR machine, and immediately transferred to a beaker containing 400 ml of water maintained at 90°C. The solution was then allowed to cool gradually until the water temperature reached approximately 35°C, enabling proper annealing of the oligonucleotides. The concentrations of the resulting double-stranded DNA templates were measured using a NanoDrop spectrophotometer. These templates were subsequently labeled using the nick translation method, as described above.

## RESULTS AND DISCUSSION

### The correct chromosome number of *Boechera calcarea*

Chromosome numbers were determined from root tips and young anthers of ten plants cultivated from seeds. All plants were confirmed to be diploid, with a chromosome count of 2*n* = 16 (**Fig. 1E-G; Suppl. Fig. 2, 3**). This finding contradicts the published record of 2n = 14 (Doudkin and Volkova, 2013; **Suppl. Fig. 1B**).

**Fig. 2.**
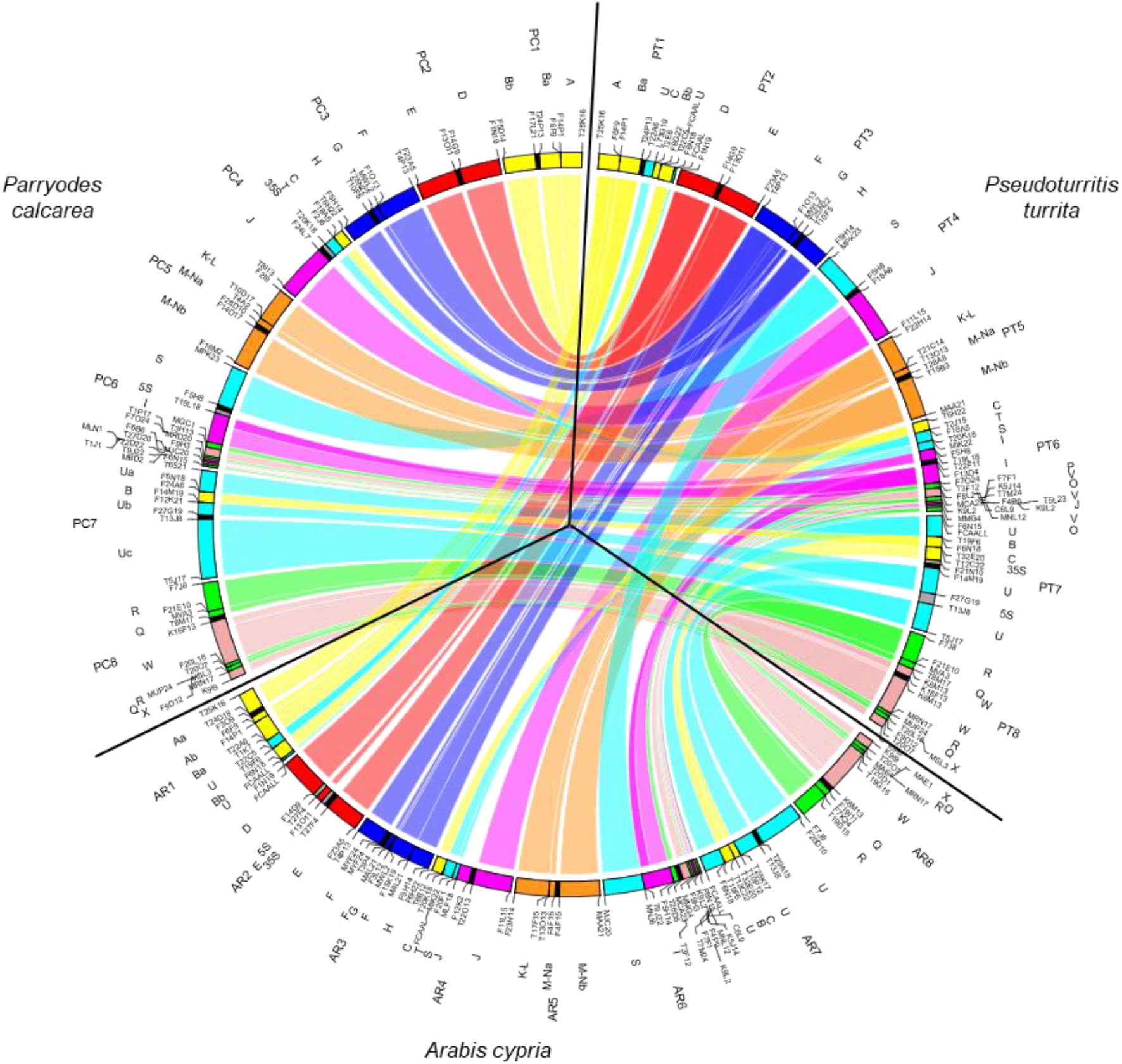
Comparative chromosome structure of *Parryodes calcarea*. A Circos diagram illustrating chromosomal collinearity between *Pseudoturritis turrita* (Mandáková et al., 2020b), *Arabis cypria* (Mandáková et al., 2020b), and *Parryodes calcarea*. Chromosomes are color-coded, with capital letters (A to X) corresponding to the eight chromosomes and 22 genome blocks (GBs) of the Ancestral Crucifer Karyotype (ACK; Mandáková et al., 2019). Black blocks represent centromeres, while grey blocks denote the locations of 35S and 5S rDNA loci. Arabidopsis BAC clones are used as markers for each (sub-)block, enabling determination of collinearity between genomes.

As all chromosome numbers in the genus *Boechera* and tribe Boechereae are based on the base number *x* = 7, Doudkin and Volkova (2013) based their taxonomic assignment of the species on morphological and karyological similarity with *Boechera* species and the fact that at least one other species of the genus is native to Russia (*B. falcata*, 2*n* = 14). However, the count of 2n = 16 provided compelling evidence that the taxon is not a Boechereae species. Considering this chromosome number and morphological characters (indumentum of simple and stalked forked trichomes, fruit a linear latiseptate silique) of the species, we hypothesized that *B. calcarea* is more likely a member of the tribe Arabideae.

### Cytogenomic analysis confirmed Arabideae-specific chromosomal structures and centromere repositioning

To test the hypothesis that the studied species belongs to the tribe Arabideae, we performed initial comparative chromosome painting on mitotic chromosomes using probes corresponding to differentially labeled BAC contigs of *Arabidopsis thaliana*. These probes were designed to represent selected combinations of genomic blocks specific to Arabideae chromosomes (Mandáková et al., 2020b): Ch1 (genomic blocks A and B), Ch4 (C and Jb), Ch7 (U) and Ch8 (R, Q, and X). This experiment clearly identified Arabideae-specific chromosomal structures in the genome of *B. calcarea* (**Suppl. Fig. 2**).

To reconstruct the entire karyotype structure of the species, we used all 22 genomic blocks designed according to the structure of the eight chromosomes of *Pseudoturritis turrita* and the younger Arabideae clades (Mandáková et al., 2020b; Nowak et al., 2020; **Fig. 2, Suppl. Fig. 2, 4**). Comparative chromosome painting revealed that the positions of genomic blocks on all eight chromosomes corresponds to those in younger Arabideae genomes (Arabideae crown group; *Arabis* and *Draba*). Perfect collinearity of genomic blocks was observed for chromosomes Pc1, Pc2, Pc3, Pc4, Pc5, and Pc8 (**Fig. 1H, 1I, 2; Suppl. Fig. 4**). On chromosome Pc7, we identified a short fragment of genomic block B within U (**Fig. 1H, I**). Our unpublished data suggest that this feature is common among all Arabideae species but was undetected in previous studies due to the lower resolution of earlier chromosome painting experiments (Mandáková et al., 2020b; Nowak et al., 2020). Chromosome Pc6 exhibited a complex reshuffling of genomic blocks, resulting in two separated parts of block O and three parts of block V (**Fig. 1H, I**). Although the structure of Pc6 slightly differs from the structure of chromosome 6 homeologs in other Arabideae genomes, we attribute these differences to the higher resolution of chromosome painting compared to earlier studies (**Fig. 2, Suppl. Fig. 4**).

Positional shift of centromeres was previously reported on five out of six chromosomes in the crown-group Arabideae (Mandáková et al., 2020b). Therefore, precise characterization of centromere positions was performed using a BAC-by-BAC approach. Chromosomes Pc1, Pc2 and Pc3 exhibited ancestral-like centromere positions as in *P. turrita*. In contrast, centromeres of chromosomes Pc5, Pc6, Pc7 and Pc8 showed independent positional shifts, as compared to homeologous centromeres in *P. turrita* as well as in crown-group Arabideae species. Structure of chromosome Pc4 aligns with that of the Arabideae crown-group species, distinguished from *P. turrita* by a reciprocal translocation. Notably, no centromere shifts were observed on this chromosome within the Arabideae crown group (**Fig. 1H, I, 2**).

Our chromosome painting data provide robust evidence for the placement of the studied species into the Arabideae. The shared chromosomal signatures with both the ancestral genome of *P. turrita* and the Arabideae crown-group species confirm that the taxon represents an intermediate transition between the ancestral and crown-group clades. Additionally, our findings reinforce previous observations that centromere repositioning serves as a predominant mechanism of chromosome differentiation in the tribe Arabideae, a phenomenon otherwise rare in the Brassicaceae (Mandáková et al., 2020b).

### Phylogenetic analysis reveals sister position of the studied species to the genus *Scapiarabis*

To gain initial insights into the phylogenetic position of the investigated species, nuclear ITS (ITS1 and ITS2) and plastid trnL-trnF regions were analyzed to construct phylogenetic trees. A preliminary large-scale phylogenetic analysis of the Brassicaceae family, based on ITS sequences using BrassiBase (data not shown), suggested the species’ placement in the tribe Arabideae. Subsequently, 349 ITS sequences from Arabideae accessions were realigned and trimmed to a final length of 616 bp. The inferred Maximum Likelihood (ML) tree (**Suppl. Fig. 5**) revealed that the studied species is sister to a clade comprising *Scapiarabis ariana* and *S. karategina*, with strong support (UFbootstrap 99%). Additionally, *Acirostrum alaschanicum* is identified as the sister species to the clade formed by *S. ariana, S. karategina*, and the studied species, with relatively strong support (UFbootstrap 84%) (**Suppl. Fig. 5**). However, the ITS analysis also indicated that the genus *Scapiarabis* is paraphyletic, as *S. saxicola* and *S. popovii* grouped with these species along with seven additional Arabideae species from different genera, forming a larger, poorly resolved clade (UFbootstrap <50%) (**Suppl. Fig. 5**).

To investigate the maternal origin of the studied species, the trnL-trnF region (trimmed alignment of 739 bp) was analyzed, including 311 Arabideae accessions. The resulting ML tree (**Suppl. Fig. 6**) showed that the studied species is sister to a monophyletic clade of all four *Scapiarabis* species (*S. ariana, S. karategina, S. popovii*, and *S. saxicola*), with a strong support (UFbootstrap 100%). The clade containing *Arcyosperma primulifolium* and *Parryodes axilliflora* was positioned basally to the *Scapiarabis* clade.

### Plastome assembly and phylogenetic analysis confirms close relationship between the studied species and *Scapiarabis saxicola*

Based on the results of the ITS and trnL-F phylogenies, we assembled the complete plastome of the studied species and incorporated all available Arabideae plastomes to construct a more robust phylogeny. The newly assembled and annotated plastome (**Suppl. Fig. 7**) has a total length of 154,218 bp, with a mean GC content of 36.3% and an average base coverage of 351.5×. In total, plastomes from 26 Arabideae accessions were included. After manual inspection and trimming, the final alignment length was 100,546 bp.

The resulting well-supported plastome phylogeny (**Fig. 3**) corroborated our earlier findings from single-marker analyses, indicating that the studied species is closely related to the genus *Scapiarabis*. Despite limited taxon sampling within *Scapiarabi*s, with only the plastome of *S. saxicola* available, the phylogeny clearly demonstrated a sister relationship between the studied species and *S. saxicola*. Furthermore, the plastome tree positioned these two species as closely related to a clade containing *Arcyosperma primulifolium* and *Parryodes axilliflora*. Pairwise alignment of the complete plastomes of the studied species and *S. saxicola* revealed a high sequence similarity of 98.8%.

**Fig. 3.**
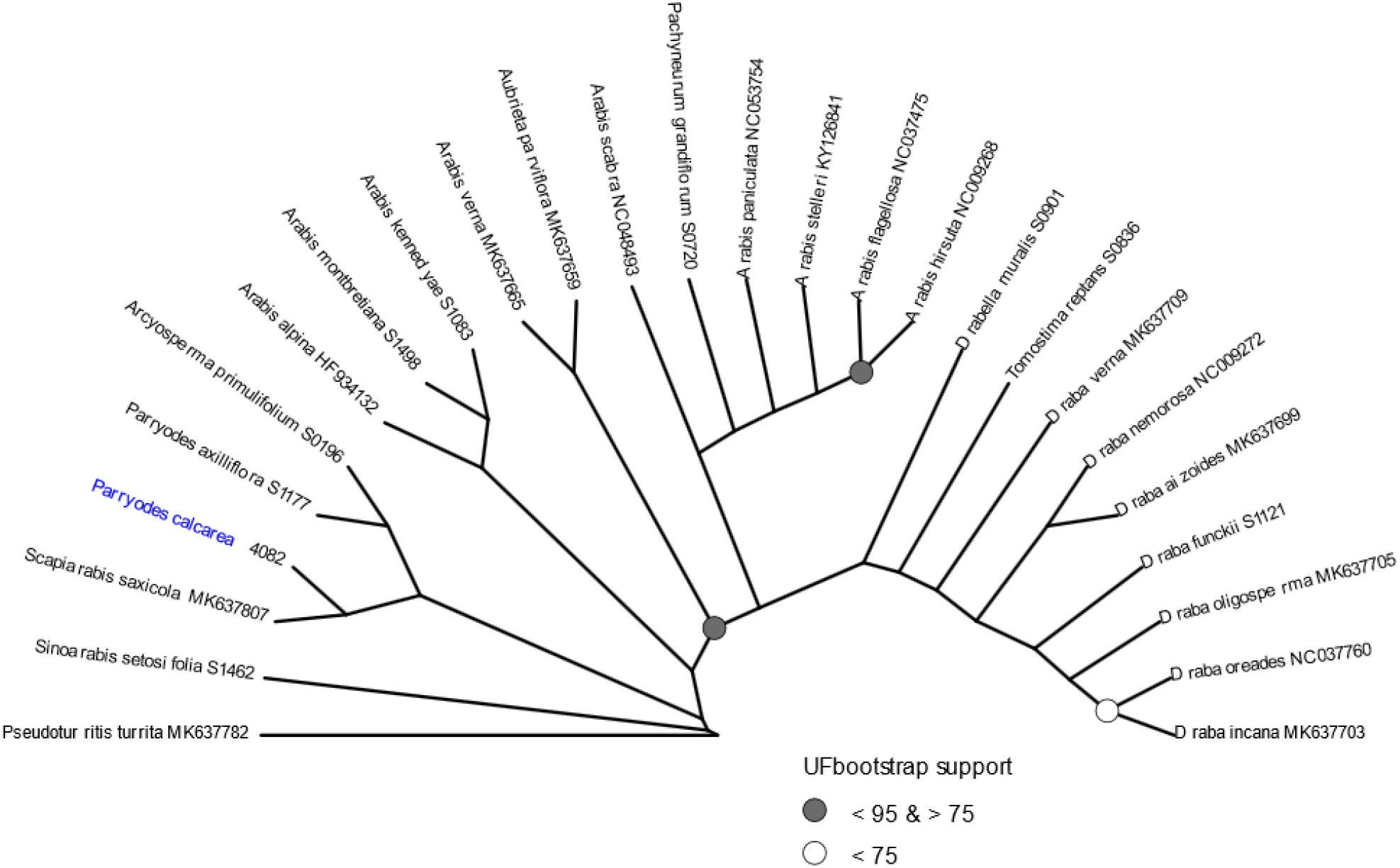
Maximum likelihood tree of the Arabideae tribe based on plastome data. Nodes with ultrafast bootstrap (UF bootstrap) support of 100% are not explicitly labeled for clarity.

Our findings align with the most complete Brassicaceae genus-level family phylogenies to date (Hendriks et al., 2023), based on nuclear (1,081 genes, 319 of the 349 genera; 57 of the 58 tribes) and plastome (60 genes, 265 genera; all tribes) data. Their ML maternal phylogeny placed *S. saxicola* as sister to the clade containing *A. primulifolium* and *Parryodes axilliflora*, consistent with our results. However, it is important to note that in the nuclear gene tree of Hendriks et al. (2023), *S. saxicola* is positioned as sister to *Acirostrum alaschanicum*, a species absent from their plastome phylogeny. Our attempts to assemble the plastome of *A. alaschanicum* from the Hyb-Seq data (SAMN31015661) used in their study were also unsuccessful due to low coverage of plastome reads.

### Repeatome analysis reveals the dominance of LTR retrotransposons and enrichment of Ty3/gypsy lineages in the studied genome

Genome size of *the* investigated species was estimated by flow cytometry to be 663,98 Mb/1C; standard deviation 6.23 (**Suppl. Fig. 8A**). Repetitive sequences were identified and characterized using the RepeatExplorer2 pipeline (Novák et al., 2020) and TAREAN (Novák et al., 2017), utilizing Illumina low-coverage data sampled to 0.5× genome coverage. Repeats were found to constitute 64.87% of the genome (**Suppl. Tab. 3**). Among these, LTR retrotransposons were the most abundant repeat type, with the Ty3/gypsy superfamily representing 49.11% of the genome. Within this superfamily, non-chromoviral Ty3/gypsy elements (Athila and Retand) were particularly dominant, accounting for 41.21%, while chromoviral lineages (CRM and Tekay) contributed 7.59% of the genome. Although the Ty1/copia superfamily comprised more lineages than Ty3/gypsy (seven vs. three), it occupied only 1.18% of the genome. Non-LTR retrotransposons, specifically LINEs, were present in low proportions (0.52%). DNA transposons made up 6.43% of the genome, represented by four main lineages: CACTA, MuDR, Harbinger, and hAT.

Tandem repeats constituted 0.40% of the genome (**Suppl. Tab. 3**). Five distinct tandem repeat sequences were identified, with monomer lengths ranging from 73 to 454 bp. The most abundant tandem repeat, Sat_105 (105 bp), accounted for 0.2% of the genome, followed by Sat_310 (310 bp; 0.09%) and Sat_177 (177 bp; 0.07%). The remaining tandem repeats, Sat_454 and Sat_73, were present at lower proportions (0.02% and 0.01%, respectively). Notably, tandem repeat Sat_177 exhibited a pairwise sequence identity of 74% with another cluster identified as a tandem repeat with a monomer size of 176 bp (0.01%; Sat_177_v2), suggesting these are likely two variants of a tandem repeat family. Clusters annotated as rDNA (5S and 35S) collectively accounted for 1.80% of the genome. Unclassified repeat clusters, which could not be assigned annotations, made up 3.10% of the genome.

When compared to other Arabideae species analyzed by Mandáková et al. (2020b), LTR retrotransposons from the Ty3/gypsy superfamily were also identified as the predominant repeat type in genome of the studied species. However, the genome displayed a notable enrichment in the Athila, Retand, and CRM lineages, which is likely responsible for its larger genome size (664 Mb). Enrichment of Ty3/gypsy LTR retroelements, particularly Athila and Retand, has also been observed in other species within the Brassicaceae (e.g., Hloušková et al., 2019; Dogan et al., 2021; Zhang et al., 2020), further supporting their significant role in genome expansion within this family.

### Shared tandem repeats reveal close genomic relationships between the studied species and *Scapiarabis saxicola*

To compare the repetitive sequences of in the genome of the studied species with those of other Arabideae species (*Acirostrum alaschanicum, Arabis auriculata, Ar. cypria, Aubrieta canescens, Parryodes axilliflora, Pseudoturritis turrita*, and *Scapiarabis saxicola*), two comparative clustering analyses were conducted using the RepeatExplorer2 pipeline. These analyses incorporated publicly available data from other Arabideae species to verify the position of the studied species within the Arabideae based on repeat profiles.

The first comparative clustering analysis, based on low-coverage Illumina data (**Suppl. Fig. 8A**), demonstrated that the genome of the studied species and *S. saxicola* have highly similar repeat profiles, with notable differences in the abundance of specific repeat types. For example, chromoviral CRM elements were more abundant in the genome of the studied species (7.30%) than in *S. saxicola* (2.35%), while tandem repeats were more prevalent in *S. saxicola*. Both species exhibited significantly higher repeat content (approximately or over 60%) compared to other Arabideae species analyzed by Mandáková et al. (2020b), whose repeat content ranged between 11.21% and 38.11% in genomes of smaller sizes (205–391 Mb). Shared LTR retroelements, particularly from the Athila and Retand lineages, are likely drivers of the increased genome size in the genome if the studied species and *S. saxicola*., the overall highly similar repeat profiles of the studied species and *S. saxicola* and the absence of species-specific repeats confirm their close phylogenetic relationship.

The second comparative clustering analysis (**Suppl.Fig. 8B**) incorporated Hyb-Seq data (Hendriks et al., 2023) alongside low-coverage Illumina sequencing data for the studied species. This analysis revealed uneven representation between the two datasets: Hyb-Seq data provided high coverage of target regions, whereas low-coverage sequencing better captured non-target regions. Despite these differences, clusters annotated as LTR retroelements (e.g., Athila, Retand, and CRM) and DNA transposons (e.g., CACTA) were consistently identified across all four genomes, with the studied genome contributing a higher proportion of reads to these clusters. Tandem repeats Sat_310 and Sat_177 were detected in all species, representing a single cluster at low abundance (0.04%), while Sat_105 was not observed. Additionally, a 444-bp satellite, shared across all genomes, was found within a supercluster representing the Athila LTR retroelement (0.94%). *Parryodes axilliflora* exhibited species-specific tandem repeats, with the most abundant being 266 bp (1.4%) and 216 bp (0.22%) in length.

The analysis further identified two tandem repeats, Sat_310 and Sat_177, shared between the studied genome, *Parryodes axilliflora*, and *Scapiarabis saxicola* (**Suppl. Fig. 9**), along with Sat_105 shared between the studied species and *S. saxicola*. Tandem repeat Sat_310 demonstrated high sequence similarity between the studied species and *S. saxicola* (97.4% pairwise identity) and between the studied species and *P. axilliflora* (91%; **Suppl. Fig. 9A**). This repeat was more abundant in *S. saxicola* (0.37%) and less so in *P. axilliflora* (0.03%) compared to the studied species. Similarly, Sat_177 exhibited pairwise identity ranging from 73% to 87.6% in *S. saxicola* (1.08%; ScSa_177_1 and ScSa_177_2) and 70.8% to 93.3% in *P. axilliflora* (0.39%; PaAx_177_1 and PaAx_177_2) compared to variants of the repeat in the studied species (**Suppl. Fig. 9B**). In contrast, Sat_105 was present in *S. saxicola* at a lower abundance (0.08%) and had a shorter consensus sequence (78 bp; **Suppl. Fig. 9C**).

Additionally, Sat_177 and its variant (Sat_177_v2) displayed sequence similarities of 87.1% and 75.7%, respectively, with tandem repeats PsTu1 and PsTu2, previously identified in *Pseudoturritis turrita* (Mandáková et al., 2020b; **Suppl. Fig. 9B**).

### Chromosome localization of tandem repeats suggests rapid centromere evolution in Arabideae

To further elucidate the karyotype structure of the studied accession, we conducted chromosome localization of idenfified tandem repeats. Arabidopsis-like telomeric repeat (TTTAGGG)_n_ and species-specific subtelomeric satellite Sat_177 were exclusively detected at the chromosome termini, confirming the absence of interstitial telomeric loci (**Fig. 1G, Suppl. Fig. 3**). In contrast, remaining four identified tandem repeats were localized within the heterochromatic pericentromeric regions. Specifically, Sat_73 and Sat_105 were detected in the pericentromere of chromosome Pc3, while the pericentromere of chromosome Pc4 contained 35S rDNA and Sat_105. Chromosome Pc6 displayed 5S rDNA and Sat_310 in its pericentromeric region, whereas Sat_310 alone was identified in the pericentromere of chromosome Pc7. The pericentromere of chromosome Pc8 harbored Sat_454 (**Fig. 1F, G, Suppl. Fig. 3**). Notably, no tandem repeats were found in the pericentromeres of chromosomes Pc1, Pc2, or Pc5. Interestingly, no centromeric satellite was shared among all chromosomes. Instead, four chromosome-specific tandem repeats were observed, suggesting rapid centromere evolution in this species. This pattern aligns with the hypothesis of accelerated centromere evolution in the tribe Arabideae (Mandáková et al., 2020b).

### Reclassification of *Boechera calcarea* as *Parryodes calcarea*

Recent evidence unequivocally demonstrates that *Boechera calcarea* does not belong to the genus *Boechera* or any other genus within the tribe Boechereae (supertribe Camelinodae; German et al., 2023). Instead, our data strongly support its placement within the tribe Arabideae (supertribe Arabodae; German et al., 2023). Within this tribe, *B. calcarea* is part of a clade of eight small genera, each comprising one to four species, which are predominantly distributed in the high mountain systems of Asia (Koch et al., 2012). Despite this broad taxonomic reassignment, the precise generic placement of *B. calcarea* remains unresolved due to phylogenetic incongruences between nuclear and chloroplast datasets, as well as discrepancies across different studies.

The nuclear ITS phylogeny supports the monophyly of the group to which *B. calcarea* belongs, yet the chloroplast trnL-F phylogeny contradicts this, presenting *B. calcarea* as a distant member of *Scapiarabis*. This incongruence complicates the taxonomic interpretation. Additionally, the nuclear ITS phylogeny renders *Scapiarabis* paraphyletic, as the clade containing *B. calcarea* excludes the type species, *S. saxicola*. If the ITS phylogeny reflects true relationships, then *S. ariana* and *S. karategina* should also be excluded from *Scapiarabis*. On the other hand, the cp phylogeny suggests that *Scapiarabis* forms a well-supported monophyletic clade sister to *B. calcarea*, further questioning its inclusion in *Scapiarabis*. Placement within *Acirostrum* also raises issues. While the ITS phylogeny suggests a potential relationship between *B. calcarea, S. ariana*, and *S. karategina*-potentially warranting their inclusion in *Acirostrum* or recognition as a new genus-the cp phylogeny does not support this. Instead, the sole member of *Acirostrum* (*A. alaschanicum*) clusters separately, and the group containing *B. calcarea* and *Scapiarabis* forms a distinct lineage.

Given these phylogenetic conflicts, an intermediate solution is required to minimize disruptions while reflecting the best-supported evolutionary relationships. Adopting a conservative approach, we follow Chepinoga et al. (2024) in reclassifying *Boechera calcarea* as *Parryodes calcarea* (Dudkin) D.A.German & Lysak. This approach draws a clear distinction between species with latiseptate fruits and accumbent cotyledons, such as *P. calcarea*, and *Arcyosperma primulifolium*, which possesses terete fruits and incumbent cotyledons and occupies a basal position in the ITS-based phylogeny. Additionally, the ITS phylogeny shows that *P. calcarea* is not nested within the existing lineages of *Scapiarabis* or *Acirostrum*, making its placement in *Parryodes* the most parsimonious solution under the current evidence.

This reclassification is further supported by its ability to minimize taxonomic disruptions among closely related genera. Including *B. calcarea* in *Parryodes* avoids the need to redefine or merge existing genera such as *Scapiarabis* or *Acirostrum*. However, it does not resolve the broader phylogenetic challenges posed by the relationships among *Parryodes, Pseudodraba, Sinoarabis, Baimashania*, and other genera within this clade. These genera form a phylogenetically remote clade in the cp phylogeny, underscoring the need for further investigation.

## CONCLUSIONS

This study reclassified *Boechera calcarea* as *Parryodes calcarea*, a distinct species of one of the more ancestral clades in the tribe Arabideae. Comprehensive cytogenetic and phylogenetic analyses revealed a diploid chromosome number of 2*n* = 16, Arabideae-specific chromosomal signatures inlcuding multiple centromere repositioning events. Our work excludes the described *B. calcarea* from the genus *Boechera* and left *B. falcata* as the sole extralimital *Boechera* species in Asia. These findings emphasize the value of integrative approaches in resolving taxonomic ambiguities and advancing our understanding of Brassicaceae diversification.

## ACKNOWLEDGEMENT

This work was supported by the Czech Science Foundation (projects 21-06839S and 24-11371S to TM, and 23-06840S to MAL), and Masaryk University Grant Agency (project MUNI/R/1268/2022 to TM). Additional support was provided as part of a long-term research project of the Czech Academy of Sciences, Institute of Botany (RVO 67985939). Core Facility Genomics of CEITEC Masaryk University is gratefully acknowledged for the obtaining of the scientific data presented in this paper. Computational resources for RepeatExplorer analysis were provided by the ELIXIR-CZ project (LM2023055), part of the international ELIXIR infrastructure. The authors are indebted to Roman V. Doudkin for kind providing us with seeds of *Parryodes calcarea*.

## SUPPLEMENTARY INFORMATION

**Suppl. Tab. 1** List of Arabideae species with assembled plastomes used in this study.

**Suppl. Tab. 2** Oligonucleotide probes specific to tandem repeats of *Parryodes calcarea*.

**Suppl. Tab. 3** Proportions of repetitive sequences in *Parryodes calcarea* genome.

**Suppl. Fig. 1** Botanical drawing **(A)** and chromosome figure **(B)** of *Boechera calcarea* from Doudkin and Volkova (2013).

**Suppl. Fig. 2** Comparative chromosome painting of mitotic chromosomes of *Parryodes calcarea*, prepared from root tips and hybridized with Arabidopsis BAC clone pools grouped into contigs representing marker genomic block combinations of Arabideae crown-group species (Mandáková et al., 2020b). Chromosomes were counterstained with DAPI. Scale bars = 10 µm.

**Suppl. Fig. 3** Chromosome localization of the selected tandem repeats on mitotic metaphase chromosomes of *Parryodes calcarea*. Chromosomes were counterstained by DAPI; FISH signals are shown in colour as indicated. Detailed information on the localized repeats is provided in **Suppl. Tab. 2**. Scale bars, 10 µm.

**Suppl. Fig. 4** Comparative chromosome structure of *Parryodes calcarea*. A Circos diagram illustrating chromosomal collinearity between *Pseudoturritis turrita* (Mandáková et al., 2020b) and *Parryodes calcarea*. Chromosomes are color-coded, with capital letters (A to X) representing the eight chromosomes and 22 genome blocks (GBs) of the Ancestral Crucifer Karyotype (ACK; Mandáková et al., 2019). Black blocks indicate centromeres, and grey blocks mark the positions of 35S and 5S rDNA loci. Arabidopsis BAC clones serve as markers for each (sub-)block, facilitating the comparison of collinearity between genomes.

**Suppl. Fig. 5** Maximum likelihood tree of the tribe Arabideae inferred from ITS1/ITS2 sequences. Nodes with ultrafast bootstrap (UF bootstrap) values below 50% are collapsed for clarity.

**Suppl. Fig. 6** Maximum likelihood tree of the tribe Arabideae inferred from plastid trnL-trnF sequences. Nodes with ultrafast bootstrap (UF bootstrap) values below 50% are collapsed for clarity.

**Suppl. Fig. 7** Newly assembled and annotated plastome of *Parryodes calcarea*.

**Suppl. Fig. 8** Comparative clustering analysis of repetitive sequences in selected species. **(A)** Genome sizes and the proportions of identified repetitive sequences for each species, based on data from Mandáková et al. (2020b) and *Parryodes calcarea*. **(B)** Analysis of *Arabis auriculata, Arabis cypria, Aubrieta canescens, Parryodes calcarea, Pseudoturritis turrita*, and *Scapiarabis saxicola*. **(C)** Analysis of *Acirostrum alaschanicum, Parryodes axilliflora, Parryodes calcarea*, and *Scapiarabis saxicola*. The sequence composition of the 200 most abundant clusters is displayed, with the size of each rectangle proportional to the genome proportion of a cluster for each species (*Acirostrum alaschanicum*: AcAl, *Arabis auriculata*: ArAu, *Arabis cypria*: ArCy, *Aubrieta canescens*: AuCa, *Parryodes axilliflora*: PaAx, *Parryodes calcarea*: PaCa, *Pseudoturritis turrita*: PsTu, *Scapiarabis saxicola*: ScSa). The bar plot in the top row indicates cluster sizes based on the number of reads in the comparative analysis.

**Suppl. Fig. 9** Multiple sequence alignments of shared tandem repeats across species. Comparisons of tandem repeats between *Parryodes calcarea* (Sat_310, Sat_177, and Sat_105), *Parryodes axilliflora* (PaAx), *Pseudoturritis turrita* (PsTu), and *Scapiarabis saxicola* (ScSa). **(A)** Alignment of Sat_310. **(B)** Alignment of Sat_177. **(C)** Alignment of Sat_105.

## Notes

### Competing Interest Statement

The authors have declared no competing interest.

## REFERENCES

Abbott, R. J., Smith, L. C., Milne, R. I., Crawford, R. M. M., Wolff, K., & Balfour, J. (2000). Molecular analysis of plant migration and refugia in the Arctic. Science, 289(5483), 1343–1346.

Al-Shehbaz, I. A. (2003). Transfer of most North American species of Arabis to Boechera (Brassicaceae). Novon, 13(4), 381–391.

Al-Shehbaz, I. A. (2005). Nomenclatural notes on Eurasian Arabis (Brassicaceae). Novon, 15(2), 278–280.

Al-Shehbaz, I. A., Beilstein, M. A., & Kellogg, E. A. (2006). Systematics and phylogeny of the Brassicaceae: An overview. Plant Systematics and Evolution, 259, 89–120.

Beck, J., Alexander, P., Allphin, L., Al-Shehbaz, I., Rushworth, C., Bailey, C. D., & Windham, M. D. (2012). Does hybridization drive the transition to asexuality in diploid Boechera (Brassicaceae)? Evolution, 66(4), 985–995.

Berkutenko, A. N. (1983). Krestotsvetnye Kolymskogo nagor’ya [Cruciferae of the Kolyma Upland]. Vladivostok: Academy of Sciences of the USSR.

Berkutenko, A. N., & Gurzenkov, N. N. (1976). Chromosome numbers in some species of the family Brassicaceae from the Russian Far East. Botanicheskii Zhurnal, 61(8), 1123–1126.

Bolger, A. M., Lohse, M., & Usadel, B. (2014). Trimmomatic: A flexible trimmer for Illumina sequence data. Bioinformatics, 30(15), 2114–2120.

Brigham-Grette, J., Elias, S. A., & Hopkins, D. M. (2013). The Bering Land Bridge. Oxford Research Encyclopedia of Environmental Science.

Busch, N. A. (1926). Cruciferae (sheets 26–32). In Flora Sibiriae et Orientis (Iss. 4, pp. 393–490). Leningrad: Ivan Fedorov State Typography.

Carman, J. C., Mateo de Arias, M., Gao, L., Zhao, X., Kowallis, B. M., Sherwood, D. A., Srivastava, M. K., Dwivedi, K. K., Price, B., Watts, L., & Windham, M. D. (2019). Apospory in addition to diplospory is common in Boechera, where it may facilitate speciation by recombination-driven apomixis-to-sex reversals. Frontiers in Plant Science, 10, 724.

Chepinoga, V. V., Barkalov, V. Y., Ebel, A. L., Knyazev, M. S., Baikov, K. S., Bobrov, A. A., … & Sennikov, A. N. (2024). Checklist of vascular plants of Asian Russia. Botanica Pacifica, 13(Special Issue), 3–310.

Chumová, Z., Záveská, E., Hloušková, P., Ponert, J., Schmidt, P.-A., Čertner, M., Mandáková, T., & Trávníček, P. (2021). Repeat proliferation and partial endoreplication jointly shape the patterns of genome size evolution in orchids. The Plant Journal, 107(2), 511–524.

Dogan, M., Pouch, M., Mandáková, T., Hloušková, P., Guo, X., Winter, P., … & Lysak, M. A. (2021). Evolution of tandem repeats is mirroring post-polyploid cladogenesis in Heliophila (Brassicaceae). Frontiers in Plant Science, 11, 607893.

Doležel, J., Greilhuber, J., & Suda, J. (2007). Estimation of nuclear DNA content in plants using flow cytometry. Nature Protocols, 2(9), 2233–2244.

Doudkin, R. V. (1998). On the flora and vegetation of the Lozovyi area of the Chandalaz Range. Botanicheskii Zhurnal, 83(11), 107–111.

Doudkin, R. V., & Volkova, S. A. (2013). A new species of Boechera (Brassicaceae) from the Primorsky Territory, Russia. Novon, 22(4), 411–414.

German, D. A., Hendriks, K. P., Koch, M. A., Lens, F., Lysak, M. A., Bailey, C. D., Mummenhoff, K., & Al-Shehbaz, I. A. (2023). An updated classification of the Brassicaceae (Cruciferae). PhytoKeys, 220, 127–144.

Greiner, S., Lehwark, P., & Bock, R. (2019). OrganellarGenomeDRAW (OGDRAW) version 1.3.1: Expanded toolkit for the graphical visualization of organellar genomes. Nucleic Acids Research, 47, W59–W64.

Guo, X., Mandáková, T., Trachtová, K., Özüdoğru, B., Liu, J., & Lysak, M. A. (2021). Linked by ancestral bonds: Multiple whole-genome duplications and reticulate evolution in a Brassicaceae tribe. Molecular Biology and Evolution, 38(7), 1695–1714.

Hendriks, K. P., Kiefer, C., Al-Shehbaz, I. A., Bailey, C. D., van Huysduynen, A. H., Nikolov, L. A., … & Lens, F. (2023). Less is more: Global Brassicaceae phylogeny based on filtering of 1,000 gene dataset. Current Biology, 33(1), 1–17.

Hloušková, P., Mandáková, T., Pouch, M., Trávníček, P., & Lysak, M. A. (2019). The large genome size variation in the Hesperis clade was shaped by the prevalent proliferation of DNA repeats and rarer genome downsizing. Annals of Botany, 124, 103–120.

Hopkins, R. C. (1937). Contributions to the taxonomy of Arabis. American Journal of Botany, 24(8), 591–600.

Ijdo, J. W., Wells, R. A. W., Baldini, A., & Reeders, S. T. (1991). The origin of human chromosome 2: An ancestral telomere-telomere fusion. Proceedings of the National Academy of Sciences of the USA, 88, 9051–9055.

Jin, J.-J., Yu, W.-B., Yang, J.-B., Song, Y., dePamphilis, C. W., Yi, T.-S., & Li, D.-Z. (2020). GetOrganelle: A fast and versatile toolkit for accurate de novo assembly of organelle genomes. Genome Biology, 21, 241.

Johnson, M. G., Pokorny, L., Dodsworth, S., Botigué, L. R., Cowan, R. S., Devault, A., … & Forest, F. (2018). A universal probe set for targeted sequencing of 353 nuclear genes from any flowering plant designed using k-medoids clustering. Systematic Biology, 68(4), 594–606.

Junier, T., & Zdobnov, E. M. (2010). The Newick utilities: High-throughput phylogenetic tree processing in the UNIX shell. Bioinformatics, 26(13), 1669–1670.

Karl, R., & Koch, M. A. (2013). A worldwide perspective on crucifer speciation and evolution: Phylogenetics, biogeography, and trait evolution in tribe Arabideae. Annals of Botany, 112(5), 983–1001.

Katoh, K., & Standley, D. M. (2013). MAFFT multiple sequence alignment software version 7: Improvements in performance and usability. Molecular Biology and Evolution, 30(4), 772–780.

Kiefer, C., Dobeš, C., & Koch, M. A. (2009). Boechera or not? Phylogeny and phylogeography of eastern North American Arabis (Brassicaceae). Taxon, 58(4), 1109–1121.

Kiefer, M., Schmickl, R., German, D. A., Mandáková, T., Lysak, M. A., Al-Shehbaz, I. A., Franzke, A., Mummenhoff, K., Stamatakis, A., & Koch, M. A. (2014). BrassiBase: Introduction to a novel knowledge database on Brassicaceae evolution. Plant and Cell Physiology, 55(e3), e3.

Koch, M. A., Haubold, B., & Mitchell-Olds, T. (1999). Molecular systematics and evolution of Arabidopsis and Arabis. Plant Biology, 1, 529–537.

Koch, M. A., Haubold, B., & Mitchell-Olds, T. (2001). Molecular systematics of the Brassicaceae: Evidence from coding plastidic matK and nuclear chs sequences. American Journal of Botany, 88(3), 534–544.

Koch, M., Dobeš, C., & Mitchell-Olds, T. (2003). Multiple hybrid formation in natural populations: Concerted evolution of the internal transcribed spacer of nuclear ribosomal DNA (ITS) in North American Arabis divaricarpa (Brassicaceae). Molecular Biology and Evolution, 20, 335–348.

Koch, M. A., Karl, R., German, D. A., & Al-Shehbaz, I. A. (2012). Systematics, taxonomy, and biogeography of three new Asian genera of Brassicaceae tribe Arabideae: An ancient distribution circle around the Asian high mountains. Taxon, 61(5), 955–969.

Krzywinski, M., Schein, J., Birol, I., Connors, J., Gascoyne, R., Horsman, D., Jones, S. J., & Marra, M. A. (2009). Circos: An information aesthetic for comparative genomics. Genome Research, 19(9), 1639–1645.

Löve, Á., & Löve, D. (1976). Cytotaxonomy of the Arctic-Alpine flora. Arctic and Alpine Research, 8(2), 127–136.

Mandáková, T., & Lysak, M. A. (2022). The identification of the missing maternal genome of the allohexaploid camelina (Camelina sativa). Plant Journal, 112(4), 622–629.

Mandáková, T., Pouch, M., Brock, J. R., Al-Shehbaz, I. A., & Lysak, M. A. (2019). Origin and evolution of diploid and allopolyploid camelina genomes was accompanied by chromosome shattering. Plant Cell, 31(10), 2596– 2612.

Mandáková, T., Price, B., Ashby, K., Hloušková, P., Windham, M. D., Mitchell-Olds, T., Carman, J., & Lysak, M. A. (2020a). The origin of the Boechereae (Brassicaceae) was preceded by descending dysploidy, whereas intratribal cladogenesis was marked by overall genome stasis. Frontiers in Plant Science, 11, 514.

Mandáková, T., & Lysak, M. A. (2016a). Chromosome preparation for cytogenetic analyses in Arabidopsis. Current Protocols in Plant Biology, 1(1), 1–9.

Mandáková, T., & Lysak, M. A. (2016b). Painting of Arabidopsis chromosomes with chromosome-specific BAC clones. Current Protocols in Plant Biology, 1(1), 43–51.

Mandáková, T., Hloušková, P., Koch, M. A., & Lysak, M. A. (2020b). Genome evolution in Arabideae was marked by frequent centromere repositioning. Plant Cell, 32(3), 650–665.

Minh, B. Q., Schmidt, H. A., Chernomor, O., Schrempf, D., Woodhams, M. D., von Haeseler, A., & Lanfear, R. (2020). IQ-TREE 2: New models and efficient methods for phylogenetic inference in the genomic era. Molecular Biology and Evolution, 37(5), 1530–1534.

Neumann, P., Novák, P., Hoštáková, N., & Macas, J. (2019). Systematic survey of plant LTR-retrotransposons elucidates phylogenetic relationships of their polyprotein domains and provides a reference for element classification. Mobile DNA, 10, 1–17.

Nikolov, L. A., Shushkov, P., Nevado, B., Gan, X., Al-Shehbaz, I. A., Filatov, D., Bailey, C. D., & Tsiantis, M. (2019). Resolving the backbone of the Brassicaceae phylogeny for investigating trait diversity. New Phytologist, 222(3), 1638–1651.

Novák, P., Ávila Robledillo, L., Koblížková, A., Vrbová, I., Neumann, P., & Macas, J. (2017). TAREAN: A computational tool for identification and characterization of satellite DNA from unassembled short reads. Nucleic Acids Research, 45(e111), e111.

Novák, P., Neumann, P., & Macas, J. (2020). Global analysis of repetitive DNA from unassembled sequence reads using RepeatExplorer2. Nature Protocols, 15(11), 3745–3776.

Nowak, M. D., Birkeland, S., Mandáková, T., Choudhury, R. R., Guo, X., Gustafsson, A. L. S., … & Brochmann, C. (2021). The genome of Draba nivalis shows signatures of adaptation to the extreme environmental stresses of the Arctic. Molecular Ecology Resources, 21(3), 661–676.

Revell, L. (2024). Phytools 2.0: An updated R ecosystem for phylogenetic comparative methods (and other things). PeerJ, 12, e16505. 10.7717/peerj.16505

Rollins, R. C. (1941). The species of Arabis in North America. Contributions from the Gray Herbarium of Harvard University, 142, 1–200.

Schulz, O. E. (1936). Cruciferae. In A. Engler & H. Harms (Eds.), Die natürlichen Pflanzenfamilien (Vol. 17B, pp. 227–658). Leipzig: Verlag von Wilhelm Englemann.

Smith, J. R., & Taylor, C. J. (2020). Morphological convergence and its impact on taxonomy. Taxon, 69(3), 520– 535.

Tillich, M., Lehwark, P., Pellizzer, T., Ulbricht-Jones, E. S., Fischer, A., Bock, R., & Greiner, S. (2017). GeSeq – Versatile and accurate annotation of organelle genomes. Nucleic Acids Research, 45(W6), W6–W11.

Warwick, S. I., & Sauder, C. A. (2005). Phylogeny of tribes Boechereae and Arabideae (Brassicaceae) based on chloroplast sequence data and nuclear ribosomal ITS region. Taxon, 54(2), 347–361.

White, T., Bruns, T., Lee, S., Taylor, J., Innis, M., Gelfand, D., & Sninsky, J. (1990). Amplification and direct sequencing of fungal ribosomal RNA genes for phylogenetics. In M. Innis, D. Gelfand, J. Sninsky, & T. White (Eds.), PCR Protocols: A Guide to Methods and Applications (pp. 315–322). Academic Press.

Windham, M. D., & Al-Shehbaz, I. A. (2006). New and noteworthy species of Boechera (Brassicaceae) II: Apomictic hybrids. Harvard Papers in Botany, 11(2), 257–274.

Windham, M. D., & Al-Shehbaz, I. A. (2007). New and noteworthy species of Boechera (Brassicaceae) I: Sexual diploids. Harvard Papers in Botany, 12(2), 335–355.

Yurtsev, B. A., & Zhukova, P. G. (1972). Chromosome numbers in some Arctic plants. Botanicheskii Zhurnal, 57(7), 899–902.

Zhang, S. J., Liu, L., Yang, R., & Wang, X. (2020). Genome size evolution mediated by Gypsy retrotransposons in Brassicaceae. Genomics, Proteomics & Bioinformatics, 18(3), 321–332.

